# Poly-dispersed-acid-functionalized-single-walled carbon nanotubes (AF-SWCNTs) are potent inhibitor of BCG induced inflammatory response in macrophage cell lines

**DOI:** 10.1101/2020.04.23.056960

**Authors:** Deepika Bhardwaj, Rajiv K. Saxena

## Abstract

Present study is focussed on the modulation of *Mycobacterium bovis* BCG induced inflammatory response by poly-dispersed acid-functionalized single-walled carbon nanotubes (AF-SWCNTs) in macrophages. Flow cytometric and confocal microscopy studies indicated that both BCG and AF-SWCNTs were efficiently internalized by RAW 264.7 and MH-S macrophage cell lines and were essentially localized in the cytoplasmic area. The results indicated strong antioxidant activity of AF-SWCNTs in mitigating BCG induced oxidative and nitrosative stress. We also found a marked decline in expression of BCG induced pro-inflammatory genes like COX-2, iNOS, TNF-α, IL-6 and IL-1β on treatment with AF-SWCNTs at transcriptional level. Decline in expression of BCG induced COX-2 by AF-SWCNTs was also confirmed at protein level using Western blotting. Anti-inflammatory activity of AF-SWCNTs was further validated by our results showing that AF-SWCNTs treatment induced a precipitous decline in BCG induced release of Matrix Metalloproteinases MMP-2 and MMP-9 by macrophage cell lines, by using Gelatin zymography. Taken together, our results demonstrate potent anti-inflammatory role of AF-SWCNTs in alleviating BCG induced inflammation.

## Introduction

Ever since their discovery by Ijima in 1991[1, 2, 3, 4], carbon nanotubes (CNTs) owing to their exceptional mechanical, thermal and electric properties, have found usage in a variety of commercial applications. While the commercial use of CNTs is increasing exponentially, no efficient mechanisms for natural degradation of CNTs have been found [5, 6, 7, 8]. Therefore, after being used, CNTs would end up dispersed in the environment jeopardizing human, animal and plant health. Pristine single walled carbon nanotubes as such exist as highly hydrophobic agglomerates insoluble in aqueous media. Functionalization of CNTs has been employed to convert them to poly-dispersed form by a number of methods including acid functionalization [9, 10, 11]. Acid functionalization introduces negatively charged carboxyl and sulfonate groups on carbon atoms in CNT walls rendering them hydrophilic and easily dispersible in aqueous media. AF-SWCNTs unlike SWCNTs interact efficiently with the cells.

Our group has been studying the interactions of AF-SWCNTs with several biological systems, and have demonstrated modulatory effects of AF-SWCNTs on activation and functioning of T cells [12, 13], B cells [13, 14], antigen presenting cells [15, 16], hematopoietic activity in bone marrow [7, 17] and lung epithelial cells [18, 15, 16]. Since AF-SWCNTs have suppressive effect on many components of the immune system, we hypothesized that AF-SWCNTs may also have a generalized anti-inflammatory effect. This hypothesis has been tested in the present study by using two macrophage cell lines (RAW 267.4 and MH-S) in which inflammatory response can be induced by *Mycobacterium bovis* BCG.

Our results demonstrate that, both autoclaved BCG and AF-SWCNTs are efficiently internalized by both macrophage cell lines. Furthermore, BCG induced oxidative and nitrosative stress inside the cells is substantially mitigated by AF-SWCNTs as indicated by decline in levels of nitric oxide and reactive oxygen species and downregulation in expression of various pro-inflammatory markers. These findings will provide essential information about the modulatory behavior of AF-SWCNTs in subsiding BCG induced inflammation in macrophages and provide a basis for testing the potential of AF-SWCNTs as an agent with anti-inflammatory properties.

## Materials and Methods

### Cells

RAW 264.7 (a murine peritoneal macrophage cell line) and MH-S (a murine lung alveolar macrophage cell line) obtained from American Type Cell Culture, were cultured in RPMI 1640 culture medium supplemented with 2 mM glutamine, 4.5g glucose per litre, 10 mM HEPES buffer pH 7.2, gentamycin (0.5mg/ml) and fetal bovine serum (10% V/V).

### Reagents and other supplies

AF-SWCNTs prepared by arc-discharge method and carboxylated by nitric acid treatment, were procured from Sigma-Aldrich (Cat No. 652490). The material was obtained in form of black powder that was >90% carbon and 5-8% metals with 1-3% carbon atoms as carboxyl acid groups. Average dimension of the nanotubes was 1.4 nm and its bundles ranged 4-5 nm in width and 500 to 1500 nm in length. The product was easily dispersible in water at the concentrations used. Zetasizer analysis indicated a zeta potential of −57.6 mV and average relative size of 142.6 nm (Online Resource 1a). Mid and near infrared spectroscopic results provided by Sigma-Aldrich are shown in Online Resource 1b. TEM picture of AF-SWCNTs revealed their fibrous structure (Online Resource 1c). N-Hydroxysuccinimide (NHS), N-Ethyl-N′-(3-dimethylaminopropyl) carbodiimide hydrochloride (EDAC), 2-(N-morpholino) ethanesulfonic acid (MES), Bio-Rad Clarity Western ECL substrate and 2′,7′-Dichlorofluorescin diacetate (DCFDA) were procured from Sigma Aldrich, USA. Alexafluor 633 hydrazide and DiI C-18 were obtained from Molecular Probes (Carlsbad, CA, USA). N-1-napthylethylenediamine dihydrochloride (NED), sulfanilamide, sodium nitrite and Polyclonal antibody against mouse COX-2 (Item no. 160126) were obtained from Cayman Chemicals, USA. SYBR-iTAQ Universal SYBR Green super mix was purchased from Bio-Rad Laboratories (California, USA). cDNA Synthesis kit and RNA Iso Plus Trizol were purchased from DSS TaKara, India. DMEM and RPMI 1640 culture medium supplemented with glutamine (2 mM), HEPES buffer pH 7.2 (10 mM), gentamycin (10 µg/ml), fetal bovine serum (FBS) (10% V/V), Middlebrook 7H9 media and OADC supplements were purchased from Himedia, India. Centricon 3kDa centrifugal filter device was obtained from Millipore (Billerica, MA, USA).

### Culture of BCG and fluorescence tagging using DiI C-18 dye

BCG was cultured in Middlebrook 7H9 liquid media supplemented with 10% OADC supplements. Growth was monitored by measuring absorbance of the bacterial suspension at 600 nm. Viability and cfu count was determined by plating on 7H11 agar OADC plates. BCG suspensions were autoclaved at 121° C and 15 psi for 60 minutes. Before use, the autoclaved BCG suspension was passed through a 26-gauge syringe to break any clumps. For tagging with fluorescent dye Dil C-18, BCG (1 × 10^8^) cfu were suspended in 500 µl of phosphate-buffered saline (PBS) containing 10 µM Dil C-18 dye and incubated in dark at 37°C for 2 hours followed by washing with PBS. Flow cytometric analysis indicated that by using this protocol, more than 85% of the BCG cells were tagged with Dil C-18 dye [19].

### Tagging of fluorochrome to AF-SWCNTs

Fluorescence tagged AF-SWCNTs (FAF-SWCNTs) were obtained by chemically tagging the fluorochrome Alexa Fluor 633 to AF-SWCNTs by using the MES buffer containing EDAC and NHS as described elsewhere [7], and were sonicated before use. The tagging efficiency of fluorophore to AF-SWCNTs was > 90% [14].

### Flow Cytometry

For studies about the uptake of DiI C-18 labelled BCG and FAF-SWCNTs, RAW 264.7 and MH-S cells were incubated in 24 well plate [0.1×10^6^/ml/well] with or without DiI C-18 labelled BCG (MOI 20:1) and 10 µg/ml of FAF-SWCNTs respectively. Then cells were harvested by trypsinization, washed with PBS and analysed on FACS Verse flow cytometer (BD Bioscience).

### Confocal Microscopy

To study the uptake and localization of the FAF-SWCNTs and DiI C-18 labelled BCG, both the RAW 264.7 and MH-S cells were cultured [0.1×10^6^/ml/well] on a coverslip in a 12 well culture plate, incubated with 10 µg/ml of FAF-SWCNTs and DiI C-18 labelled BCG (MOI 20:1) for 12 hours, washed twice with PBS and fixed with 4% paraformaldehyde. Nuclei were counterstained with Hoechst 33342 dye as described before [14]. Coverslips were mounted onto the glass slide with 50% glycerol and examined on a Nikon A1R Confocal Laser Scanning Microscope.

### Estimation of reactive oxygen species (ROS)

For flow cytometry studies to detect the production of ROS, both the RAW 264.7 and MH-S cells were cultured [0.1×10^6^/ml/well] in each well of 12 well plate and treated with 100 µg/ml AF-SWCNTs, Autoclaved BCG MOI 100:1 and Autoclaved BCG MOI 100:1+100 µg/ml AF-SWCNTs for 12 hours. The cells were trypsinized, washed twice with PBS, and pellet was incubated in fresh media containing 5 µM DCFDA in fresh media for 30 minutes in the dark at 37 °C. Flow cytometric analysis was performed using FACS Verse flow cytometer (BD Biosciences).

### Nitric oxide measurement

To measure the amount of Nitric oxide (NO) produced, both RAW 264.7 and MH-S cells were cultured (0.1×10^6^/ml/well in 24 well cell culture plate) in phenol red free RPMI+10% FBS and left for overnight incubation in CO_2_ incubator at 37°C and 5% CO_2_. Thereafter, cells were treated with or without 100 µg/ml AF-SWCNTs and/or Autoclaved BCG MOI 100:1 for 4 hours. NO production was measured by determining the nitrite levels in cell culture supernatants using Griess reagent as mentioned elsewhere [18]. Culture supernatants were mixed 1:1 (v/v) with Griess reagent and the color generated after 15-minute incubation at room temperature was read at 540 nm.

### Real time PCR (RT-PCR)

RAW 264.7 and MH-S cells (0.5×10^6^/ml/well) were cultured in a 6 well plate and treated with or without 100 µg/ml AF-SWCNTs and/or Autoclaved BCG MOI 100:1 for 4 hours. Following incubation, the cells were washed with cold PBS and incubated with the TRIzol reagent to extract total RNA. cDNA was synthesized using 2 µg total RNA with a cDNA synthesis kit. Forward and reverse sequences of primers were as follows:

β-actin [GGAGGGGGTTGAGGTGTT, GTGTGCACTTTTATTGGTCTCAA]

COX-2 [TGAGTGGGGTGATGAGCAAC, TTCAGAGGCAATGCGGTTCT]

iNOS [GGCAGCCTGTGAGACCTTTG, GCATTGGAAGTGAAGCGTTTC]

TNF-α [GACAAGCCTGTAGCCCACG, TGTCTTTGAGATCCATGCCGT]

IL-6 [ACAAAGCCAGAGTCCTTCAGAG, GAGCATTGGAAATTGGGGTAGG]

IL-1β [ATGCCACCTTTTGACAGTGATG, TGTGCTGCTGCGAGATTTGA]

Reaction volume of 10 µl was used for the amplification consisting of 2.5 µl of ten-fold diluted cDNA, 5 µl SYBR green, 0.2 µl (10 mM) of each primer and sterile distilled water. Cycling condition was as follows: 10 min at 95 °C and 40 cycles of 15 sec at 95 °C, 30 sec at 60 °C and the melt curve with single reaction cycle with the following conditions: 95 °C for 15 sec, 60 °C for 1 min and dissociation at 95 °C for 15 sec. After the amplification, the cycle threshold (Ct) values were obtained and normalized to the value of housekeeping gene β-actin. The relative expressions of the target genes were calculated using the 2-∆∆Ct method.

### Western Blotting

Western blot analysis was carried out as described before [20]. Briefly, RAW 264.7 and MH-S cells were cultured with or without 100 µg/ml of AF-SWCNTs and/or Autoclaved BCG MOI 100:1 for 24 hours. Cells were washed twice with ice-cold PBS and incubated in RIPA lysis buffer containing a cocktail protease inhibitor mixture at 4°C for 20 minutes and then stored at −80 degrees overnight. Next day, cell lysates were collected, sonicated 3 three times for 10 sec each and analysed for protein content using the BCA protein assay kit (Thermo Fisher scientific). Samples containing 30 µg/µl of cell lysate proteins per lane were resolved on SDS-PAGE (10% sodium dodecyl sulfate-polyacrylamide gel electrophoresis) along with pre-stained protein ladder and transferred onto PVDF membranes (Invitrogen). The transferred membranes were blocked for 2 hours using 5% BSA in TBST (25 mm Tris-HCl, pH 7.4, 125 mm NaCl, 0.05% Tween 20) and incubated with COX-2 primary antibody at 4°C overnight. Membranes were washed thrice with TBST for 5 minutes and incubated with horseradish peroxidase-coupled isotype-specific secondary antibody for 1 hour at room temperature. The immune complexes were detected by enhanced chemi-luminescence detection system (Bio-Rad) and quantified using NIH Image J software.

### Gelatin Zymography for MMP-2 and MMP-9

Protocol used for zymography has been described elsewhere [21]. Briefly, 0.1×10^6^ RAW 264.7 and MH-S cells were cultured in a 12 well cell culture plate with or without 100 µg/ml AF-SWCNTs and/or Autoclaved BCG MOI 100:1 for 12 hours in serum-free media. Supernatants were mixed with 5X non-reducing sample buffer and subjected to electrophoresis on 7.5% SDS-PAGE containing 4mg/ml gelatin. After washing with the washing buffer for 30 minutes, the gel was incubated in the incubation buffer at 37 °C for 24 hours. On the next day the gel was stained with staining solution and was destained with the destaining solution. The gelatinolytic activity of samples was detected as clear band against blue background. The bands were visualized using enhanced chemi-luminescence detection system (Bio-Rad) and quantified using NIH Image J software.

### Statistical Analysis

All experiments were repeated at least three times. Results are expressed as Mean ±SEM. The paired t-test was performed to define the significance of all the experiments. Statistical analyses were performed using Sigma Plot software (Systat software, San Jose, CA).

## Results

### Uptake of BCG and AF-SWCNTs by RAW 264.7 and MH-S cells

Uptake of BCG and AF-SWCNTs by the macrophage cell lines was examined by using Dil C-18 stained autoclaved BCG and AF-SWCNTs tagged with the fluorescent dye Alexa Fluor 633 (FAF-SWCNTs). The cells were incubated with fluorescent labelled BCG or FAF-SWCNTs for up to 24 hours and uptake of both the agents by RAW 264.7 and MH-S cells was analysed using flow cytometry and confocal microscopy. Flow cytometry results in Fig. 1A show the kinetics of uptake of labelled BCG by RAW 264.7 and MH-S cells and indicate that at 24 hour time point, almost all the cells were positive for DiI C-18 labelled BCG uptake. For confocal microscopy, cells treated with DiI C-18 labelled BCG for 12 hours were counterstained using Hoechst 33342 dye to visualize blue stained nuclei while DiI C-18 labelled BCG could be visualized by its red fluorescence. Results in Fig. 1A and Z sectioning image (3 µm depth) in the left panel of Online Resource 2 show that red fluorescence of DiI C-18 labelled BCG was localized in the cytoplasmic area of both RAW 264.7 and MH-S cells and not merely restricted to cell surface.

**Fig. 1a.**
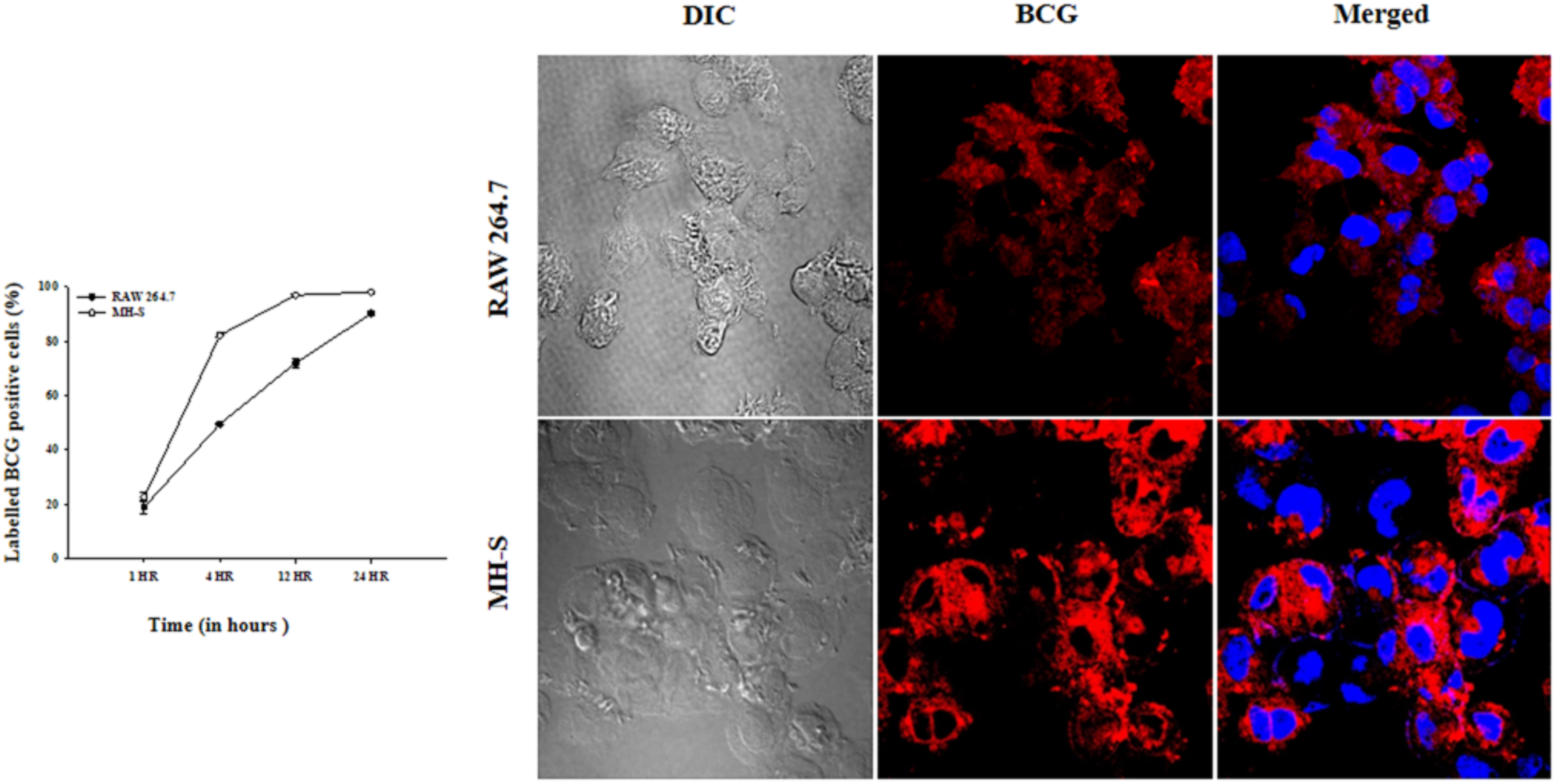
Uptake and localization of DiI C-18 labelled BCG by RAW 264.7 and MH-S cells. RAW 264.7 and MH-S cells [0.1×10^6^/ml/well] were cultured with DiI C-18 labelled BCG (MOI 20:1) for up to 24 hours. After incubation, cells were washed with PBS, detached using trypsin and analyzed on a flow cytometer to evaluate the uptake of DiI C-18 labelled BCG by RAW 264.7 and MH-S cells. Graph on the left side shows the kinetics of uptake of DiI C-18 labelled BCG where Y-axis shows labelled BCG positive cells (%). Open circles represent labelled BCG positive cells (%) for RAW 264.7 and closed circles represent labelled BCG positive cells (%) for MH-S. Each plotted value is a Mean±SEM of 3 individual experiments. Confocal microscopy results showing internalization of labelled BCG by RAW 264.7 (top panel) and MH-S (bottom panel) for 12 hour time point are shown on the right side. In each panel, micrographs show DiI C-18 labelled BCG uptake (red fluorescence) and nuclei are defined by blue stain of Hoechst 33342 dye. Magnification 100X in all cases.

Results in Fig. 1B show the kinetics of uptake of FAF-SWCNTs by RAW 264.7 and MH-S cells and indicate that at 12 and 24 hour time points almost 70%-80% cells were positive for FAF-SWCNTs uptake. For confocal microscopy, cells treated with FAF-SWCNTs for 12 hours were counterstained by Hoechst 33342 dye to visualize blue stained nuclei while FAF-SWCNTs could be visualized by their red fluorescence. Results in Fig. 1B and Z-sectioning image (3 µm depth) in the right panel of Online Resource 2 show that the red fluorescence of FAF-SWCNTs was essentially localized in the cytoplasmic area of the RAW 264.7 and MH-S cells and not merely confined to cell surface.

**Fig. 1b.**
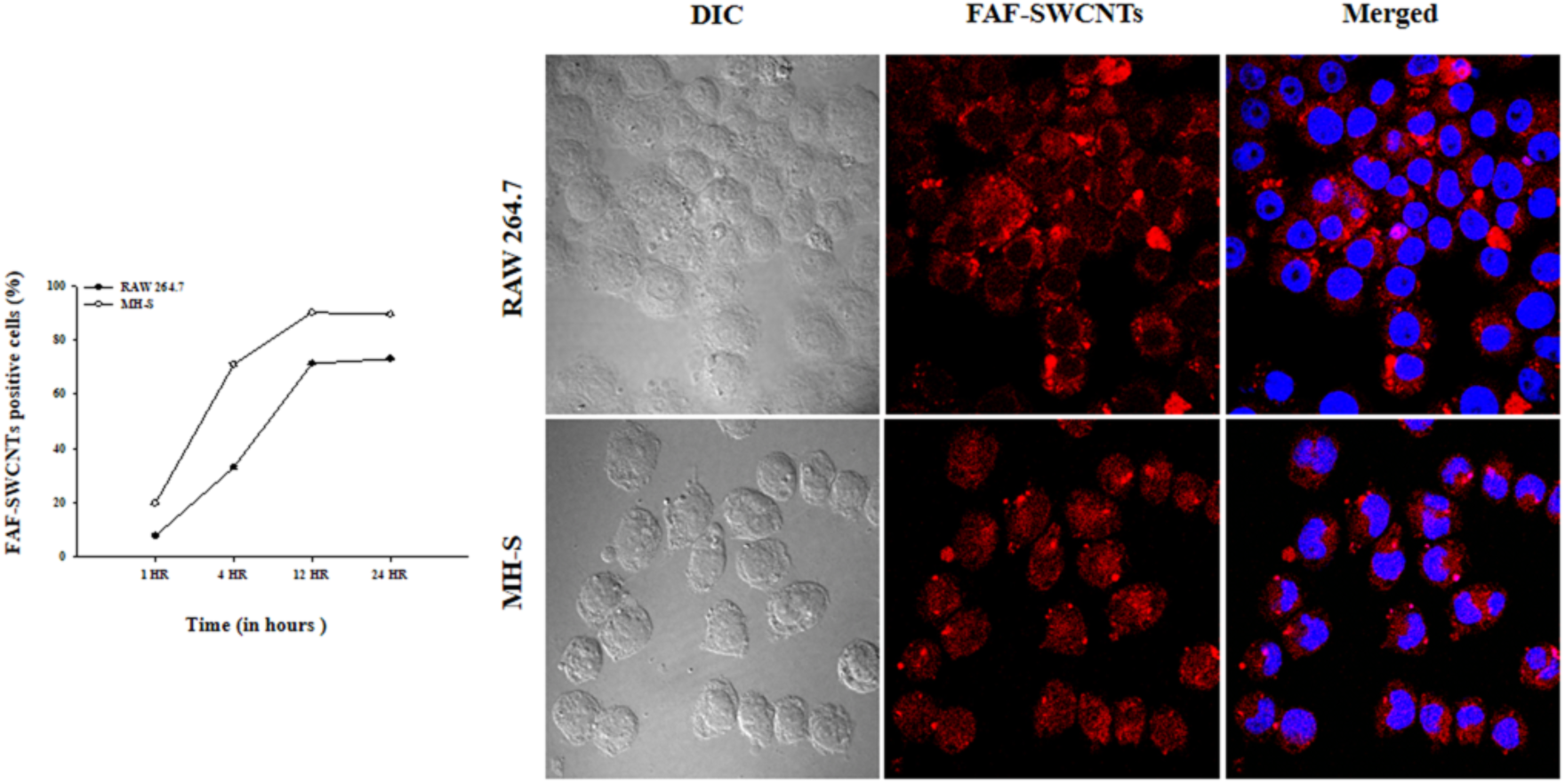
Uptake and localization of FAF-SWCNTs by RAW 264.7 and MH-S cells. RAW 264.7 and MH-S cells [0.1×10^6^/ml/well] were cultured with FAF-SWCNTs (10 µg/ml) for up to 24 hours. After incubation, cells were washed with PBS, detached using trypsin and analyzed on a flow cytometer to evaluate the uptake of FAF-SWCNTs by RAW 264.7 and MH-S cells. Graph on the left side shows the kinetics of uptake of FAF-SWCNTs where Y-axis shows FAF-SWCNTs positive cells (%). Open circles represent FAF-SWCNTs positive cells (%) for RAW 264.7 and closed circles represent FAF-SWCNTs positive cells (%) for MH-S. Each plotted value is a Mean±SEM of 3 individual experiments. Confocal microscopy results showing internalization of FAF-SWCNTs by RAW 264.7 (top panel) and MH-S (bottom panel) for 12 hour time point are shown on the right side. In each panel, micrographs show FAF-SWCNTs uptake (red fluorescence) and nuclei are defined by blue stain of Hoechst 33342 dye. Magnification 100X in all cases.

These results indicate that in both the macrophage cell lines, BCG and AF-SWCNTs get rapidly internalized and accumulate predominantly in cytoplasm. However, MH-S cells internalized higher amount of both labelled BCG and FAF-SWCNTs in comparison to RAW 264.7 cells as indicated by higher number of percent positive cells for both labelled BCG as well as FAF-SWCNTs.

### Induction of ROS production by BCG and its inhibition by AF-SWCNTs

Effect of AF-SWCNTs on BCG induced ROS production by the macrophage cell lines was examined. Results presented in Fig. 2a and 2b indicate that the untreated control cells as well as AF-SWCNTs treated cells generated little ROS. Treatment with BCG resulted in a robust ROS response that was significantly inhibited (50% and 37% inhibition in RAW 264.7 and MH-S cells respectively) in presence of AF-SWCNTs (Figure 2). These results show a potent anti-oxidant activity of AF-SWCNTs.

**Fig. 2.**
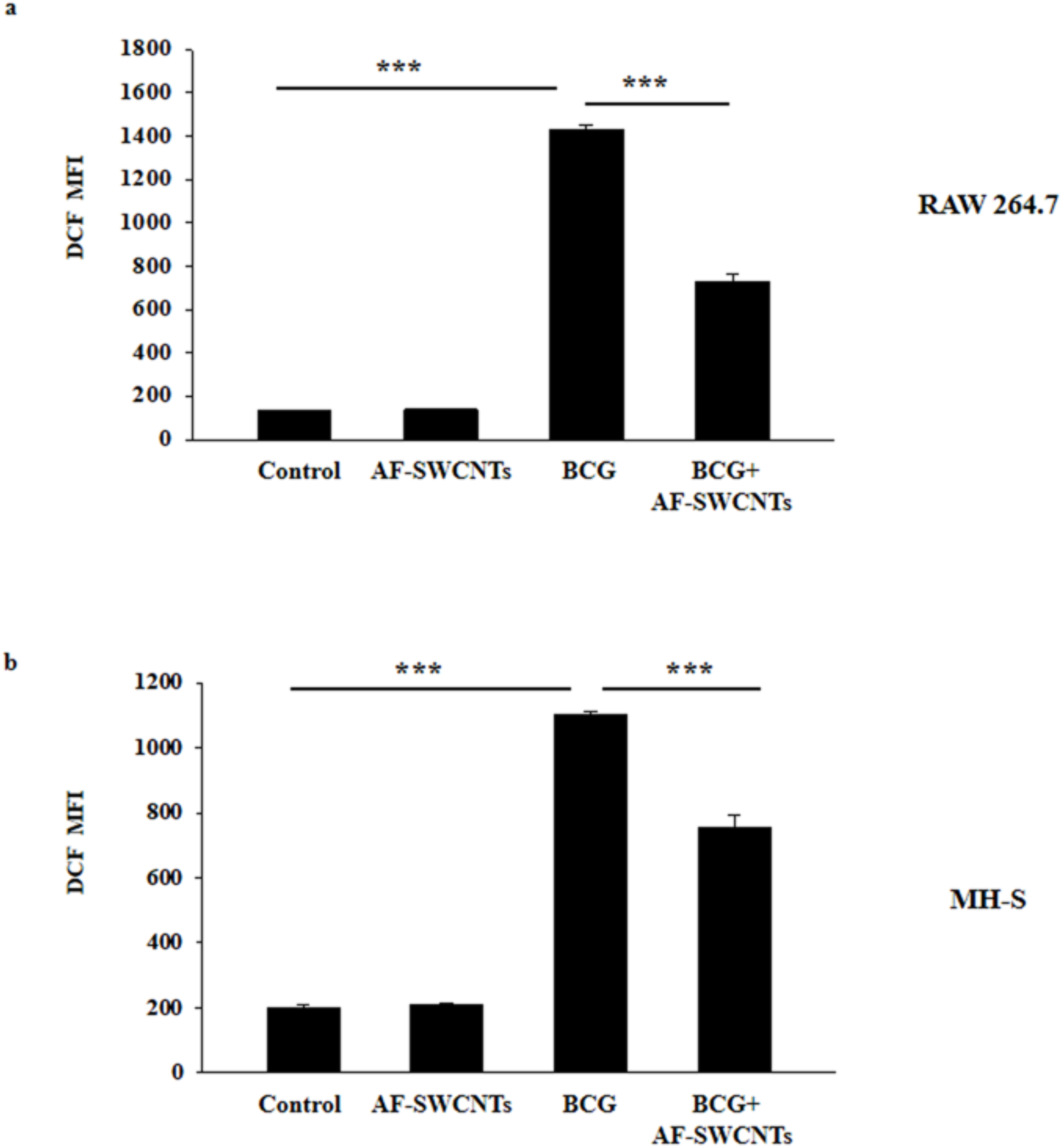
Inhibition of BCG induced ROS production by AF-SWCNTs using flow cytometry. RAW 264.7 and MH-S cells [0.1×10^6^/ml/well] were cultured with 100 µg/ml AF-SWCNTs, Autoclaved BCG MOI 100:1 and Autoclaved BCG MOI 100:1+100 µg/ml AF-SWCNTs for 12 hours. After incubation, cells were harvested, stained with 5 µM DCFDA for 30 minutes, washed to remove excess stain and acquired using flow cytometer. Panel a and b represent DCF MFI for RAW 264.7 and MH-S cells respectively. Histograms show result of 3 independent experiments (Mean ±SEM). *p<0.05, **p<0.01, ***p<0.001.

### Induction of nitric oxide production by BCG and its inhibition by AF-SWCNTs

Production of NO by macrophages on treatment with AF-SWCNTs and Autoclaved BCG was assessed by using Griess reagent. Results in Fig. 3a and 3b show that treatment with AF-SWCNTs alone did not cause any significant alteration in NO production in both the macrophage cell lines at 4 hour and 12 hour time points. However, treatment with autoclaved BCG led to a remarkable increase in NO production in RAW and MH-S cells (p<0.001) which was substantially abrogated upon treatment with AF-SWCNTs in both the cell lines; the decline being 75% in 4 hour and 12 hour time points in RAW 264.7 and 90% in MH-S in both the time points (p<0.001).

**Fig. 3.**
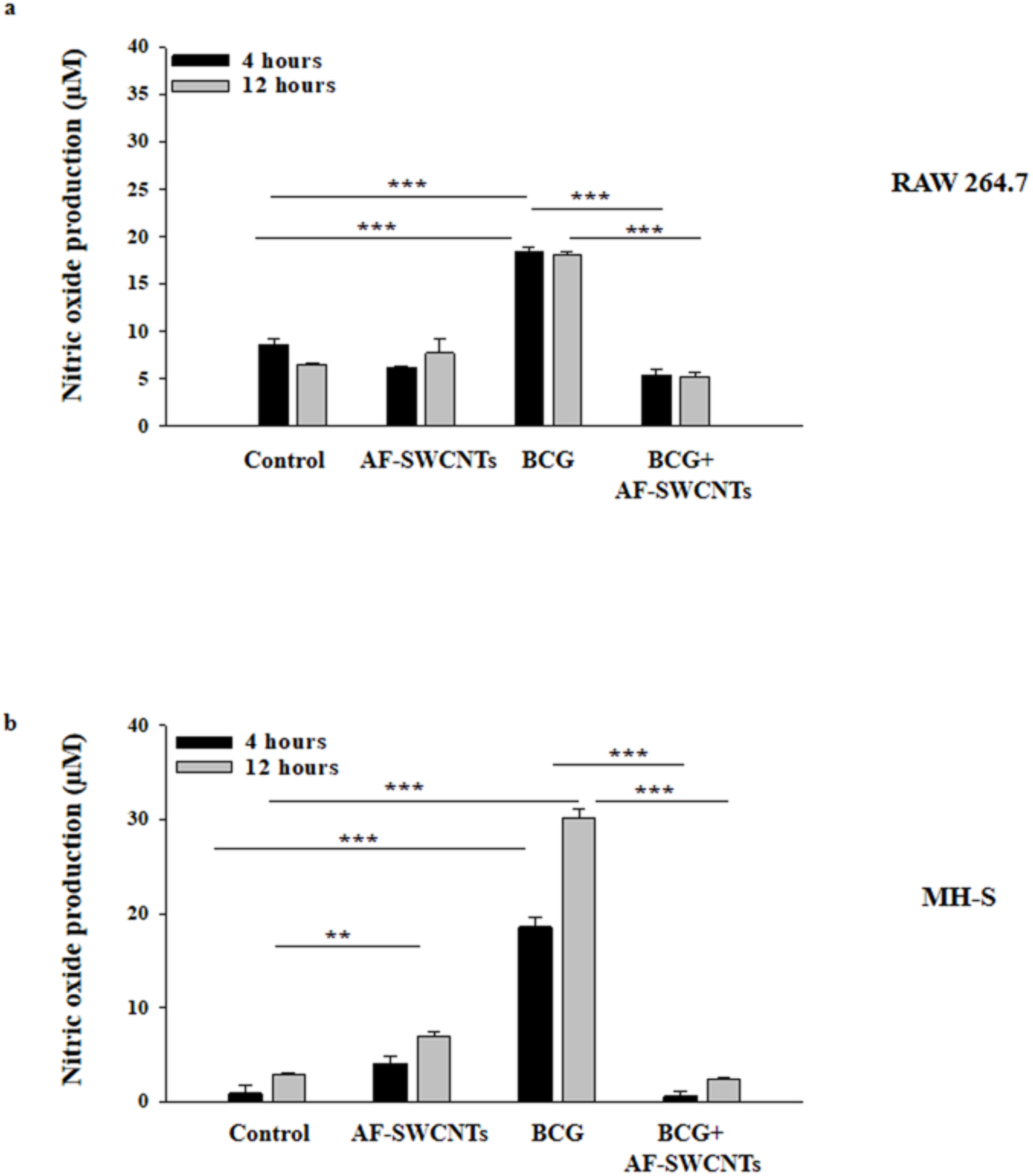
Suppression of BCG induced NO production by AF-SWCNTs by Griess assay. RAW 264.7 and MH-S cells [0.1×10^6^/ml/well] were cultured with 100 µg/ml AF-SWCNTs, Autoclaved BCG MOI 100:1 and Autoclaved BCG MOI 100:1+100 µg/ml AF-SWCNTs for 4 hours and 12 hours respectively. NO production was measured by determining the nitrite levels in cell culture supernatants using modified Griess reagent and the color generated after 30 minute incubation at room temperature was read at 540 nm using spectrophotometer. Panel a and b show nitric oxide concentration (µM) under different conditions in both 4 hour and 12 hour time points for RAW 264.7 and MH-S cells respectively where black coloured bars represent 4 hour time point and gray coloured bars represent 12 hour time point. Histograms show result of 3 independent experiments (Mean±SEM). *p<0.05, **p<0.01, ***p<0.001.

### Modulation of expression of BCG induced pro-inflammatory genes in macrophages by AF-SWCNTs

Since substantial inhibitory effect of AF-SWCNTs on BCG induced ROS and NO response was seen, we examined the expression levels of some important pro-inflammatory genes namely COX-2, iNOS, TNF-α, IL-6 and IL-1β in cells treated with BCG in presence and absence of AF-SWCNTs. Real time PCR results in Fig. 4 show that COX-2, iNOS, TNF-α, IL-6 and IL-1β were significantly upregulated in both RAW 264.7 and MH-S cells on treatment with autoclaved BCG (p<0.001). Addition of AF-SWCNTs led to a steep decline in the expression of these pro-inflammatory genes; the decline being 49% in COX-2 (p<0.001), 95% in iNOS (p<0.05), TNF-α (p<0.001) and IL-6 (p<0.001) and 63% in IL-1β (p<0.05) in RAW 264.7 while reduction of 75% in COX-2 (p<0.001), 95% in iNOS (p<0.001), 68% in TNF-α (p<0.001), 90% in IL-6 (p<0.05) and 80% in IL-1β (p<0.001) was observed in MH-S.

**Fig. 4.**
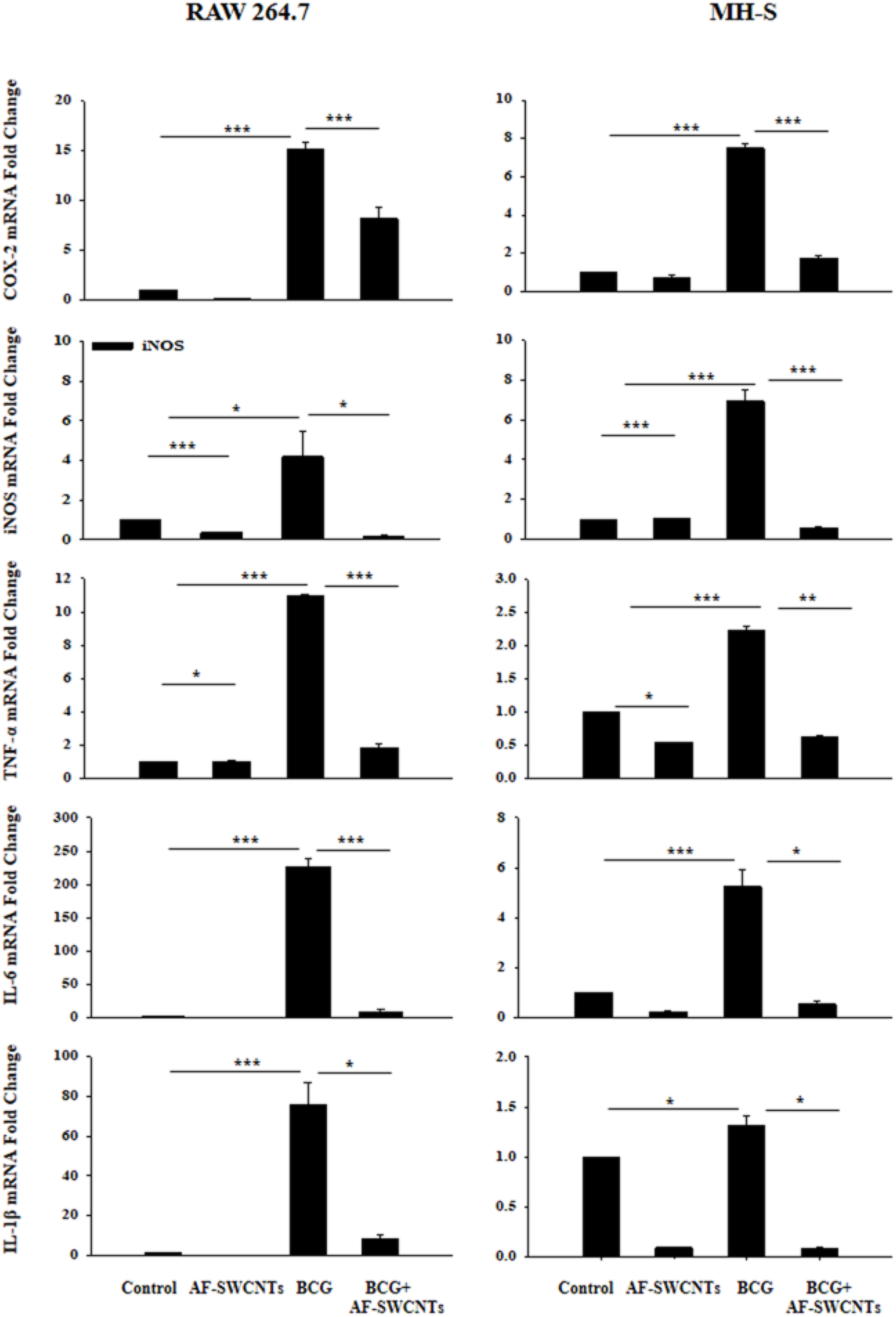
Modulation of expression of BCG induced pro-inflammatory genes in macrophages by AF-SWCNTs. RAW 264.7 and MH-S cells [0.5×10^6^/ml/well] were cultured with 100 µg/ml AF-SWCNTs, Autoclaved BCG MOI 100:1 and Autoclaved BCG MOI 100:1+100 µg/ml AF-SWCNTs for 4 hours. RNA was extracted using Trizol RNA extraction method, quantified using nanodrop and 2µg of RNA was reverse transcribed to form cDNA. Thereafter, real time PCR was performed to check relative gene expression of pro inflammatory genes under different conditions. Fold change in gene expression was calculated using 2−∆∆Ct method. Histograms show result of 3 independent experiments (Mean± SEM). *p<0.05, **p<0.01, ***p<0.001.

### Modulation of expression of BCG induced COX-2 by AF-SWCNTs using Western blotting

Expression of COX-2 in cells treated with BCG and AF-SWCNTs was also examined at protein level. Results in Fig. 5a and 5b demonstrate a substantial increase in activity of COX-2 on treatment with autoclaved BCG, which was significantly abrogated on treatment with AF-SWCNTs; the decline being 21% (p<0.05) and 85% (p<0.05) in RAW 264.7 and MH-S cells respectively.

**Fig. 5.**
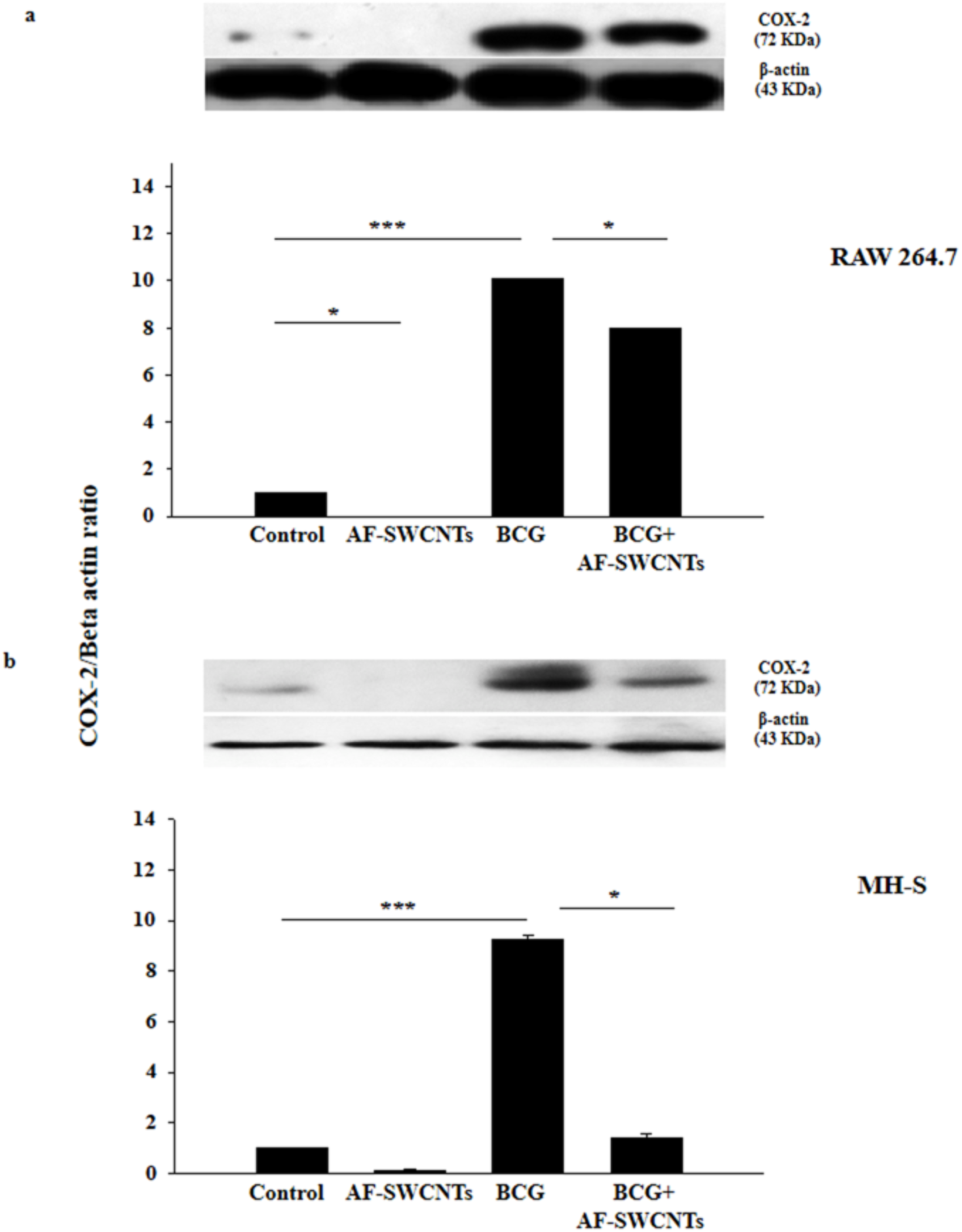
Modulation of expression of BCG induced COX-2 by AF-SWCNTs using Western blotting. RAW 264.7 and MH-S cells [0.5 × 10^6^/ml/well] were cultured with 100 µg/ml AF-SWCNTs, Autoclaved BCG MOI 100:1 and Autoclaved BCG MOI 100:1+100 µg/ml AF-SWCNTs for 24 hours. Protein lysates were prepared by using RIPA extraction buffer. The estimation of protein was done by BCA method. The samples were resolved with 10% SDS-PAGE, transferred onto PVDF membrane, nonspecific binding was blocked with 5% BSA and then probed with COX-2 antibody overnight at 4 degree Celsius. The immune-complexes were detected using HRP conjugated secondary antibody and quantified by Image-J software. Panel a and Panel b show representative blots of COX-2 normalized with housekeeping gene Beta actin along with histograms for RAW 264.7 and MH-S cells respectively. Histograms show result of 3 independent experiments (Mean± SEM). *p<0.05, **p<0.01, ***p<0.001.

### Modulation of expression of BCG induced MMP-2 and MMP-9 by AF-SWCNTs using Gelatin zymography

Matrix metalloproteinases (MMPs) are important participants in inflammatory responses. We found a significant increase in the activity of MMP-2 and MMP-9 in culture supernatants of the two cell lines treated with BCG as can be seen by a clear thick band on gelatin zymogram. However, treatment with combination of autoclaved BCG and AF-SWCNTs led to a marked decline in the activity of both MMP-2 and MMP-9 in both cell lines. Results in Fig. 6 show that treatment with autoclaved BCG led to a marked increase in activity of MMP-2 and MMP-9 in both RAW 264.7 and MH-S cells as indicated by very bright bands on the gelatin zymogram. the decline being 83% (p<0.001) in MMP-2, 64% (p<0.001) in MMP-9 in RAW 264.7 and 80% (p<0.01)in MMP-2 and 83% (p<0.001) in MMP-9 in MH-S cells.

**Fig. 6.**
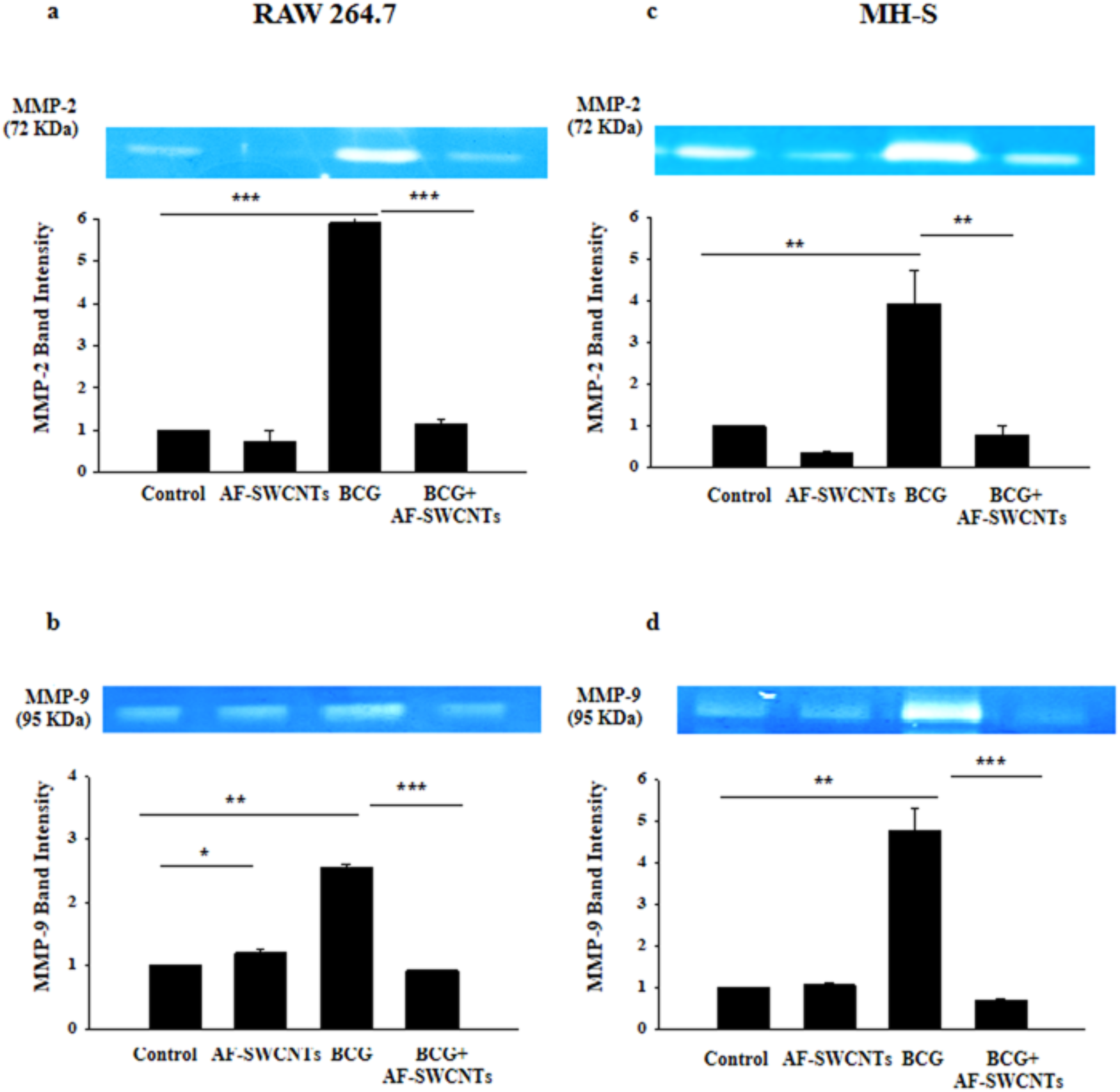
Modulation of expression of BCG induced MMP-2 and MMP-9 by AF-SWCNTs using Gelatin Zymography. RAW 264.7 and MH-S cells [0.5 × 10^6^/ml/well] were cultured with 100 µg/ml AF-SWCNTs, Autoclaved BCG MOI 100:1 and Autoclaved BCG MOI 100:1+100 µg/ml AF-SWCNTs for 12 hours in serum free medium. Thereafter, gelatin zymography was performed using cultured supernatant to check the activity of different MMPs. Panel a and b show representative gel images of MMP-2 and MMP-9 along with histograms under different conditions for RAW 264.7 cells. While Panel c and d show representative gel images of MMP-2 and MMP-9 along with histograms under different conditions for MH-S cells. Histograms show result of 3 independent experiments (Mean ± SEM). *p<0.05, **p<0.01, ***p<0.001.

## Discussion

Inflammatory responses occur in response to pathogen associated molecular patterns (PAMPs) and damage associated molecular patterns (DAMPs). In due course of time if the inducer of inflammation is lost, inflammation subsides and resolution phase takes over to repair the damage caused by inflammation. In both inducer and resolution phases, macrophages play a crucial role [22]. Regulation of inflammatory responses, especially its suppression is of great interest clinically. There are a large number of drugs in use that have anti-inflammatory properties and search is on for new candidates with their own special properties and potential for use under specific conditions.

Our previous work on AF-SWCNTs has shown that it suppresses B, T and NK cell function and processing and presentation of antigens by macrophages as well as lung epithelial cells [14, 12, 13, 16, 15, 10]. These studies therefore may suggest that AF-SWCNTs may be a good candidate as a general immunosuppressive agents and perhaps as anti-inflammatory agents. In the present study, we essentially induced inflammatory responses in two macrophage cell lines by treating with autoclaved BCG, and tested the efficacy of AF-SWCNTs to modulate these responses. Two macrophage cell lines were selected, one (RAW 264.7), a murine peritoneal macrophage cell line and other (MH-S), a mouse alveolar macrophage cell line. Both cell lines have been extensively used as models for studying macrophage functions and inflammatory responses [23, 24]. Further both cells lines effectively interacted with BCG as well as AF-SWCNTs since both were internalized rapidly by the macrophage cell lines.

Induction of inflammatory response by BCG proceeds through many steps. Entry of BCG into the cells is accompanied by mitochondrial damage, which leads to release of ROS inside the cell [25] and abrupt increase in expression of iNOS (inducible Nitric oxide synthase) enzyme [26] which brings about conversion of L-arginine to L-citrulline and NO. ROS such as superoxide combines with NO to form RNS (reactive nitrogen species), which further amplify the pro-inflammatory burden of ROS [27]. Thereafter, ROS and NO bring about formation of enzyme COX-2 from arachidonic acid. COX-2 contributes to formation of PGE2, which further exacerbates the inflammatory response [28]. DNA damage induced by ROS and NO leads to increase in transcription of pro-inflammatory genes like COX-2, iNOS, TNF-α, IL-6, IL-1β, and MMPs leading to establishment of an inflammatory condition inside the cell.

Many of the PAMP receptors (TLR molecules) are present intracellularly and internalization of BCG would ensure activation of TLR receptors leading to inflammatory response. Interestingly, AF-SWCNTs though also internalized, did not induce inflammatory response on their own indicating that AF-SWCNTs may not activate pro-inflammatory receptors inside the cell. A recent report has demonstrated the activation of inflammatory response by pristine SWCNTs and suggests a role of TLR2 and TLR4 receptors in this response [29]. Pristine SWCNTs were used in this study that are strongly hydrophobic and the interactions between SWCNTs and TLR receptors was ascribed to hydrophobic interactions between SWCNTs and TLR receptors [29]. In our study, acid functionalized SWCNTs were used that are hydrophilic in nature and may not interact with TLR receptors through hydrophobic interactions. Accordingly, in our study, AF-SWCNTs by itself did not cause significant inflammatory reaction.

Sequence of events enumerated above leading to inflammatory response in BCG exposed macrophages shows that the magnitude of the response can be assessed by measuring the levels of several participating molecules. These include reactive oxygen species [30], nitric oxide [31], matrix metalloproteinases [32] and expression of key pro-inflammatory genes like COX-2 [33], iNOS [34], TNF-α [35], IL-6 [36] and IL-1β [37] etc. We used all these parameters on the two macrophage cell lines and found that all these parameters were strongly upregulated by the BCG, a known potent inducer of inflammation. Further in all cases and for both cell lines, the upregulated inflammation parameters were substantially down regulated by AF-SWCNTs. Even though we have shown downregulation of several mediators of inflammation in response to AF-SWCNTs, how exactly AF-SWCNTs modulate the inflammatory response is not clear and will require further work.

Our results point to the possibility of use of AF-SWCNTs as an anti-inflammatory agent under certain defined conditions. Conditions indeed have to be worked out. Different doses and times have to be examined *in vitro* and *in vivo*. Additional work will also be needed to pinpoint the points in the intra-cellular pathways leading to inflammation, where AF-SWCNTs exert the suppressive influence.

## Acknowledgements

Research funding from the Department of Science and Technology, Government of India, and fellowship support to DB from DBT are gratefully acknowledged.

---

**Supplementary Fig. 1.**
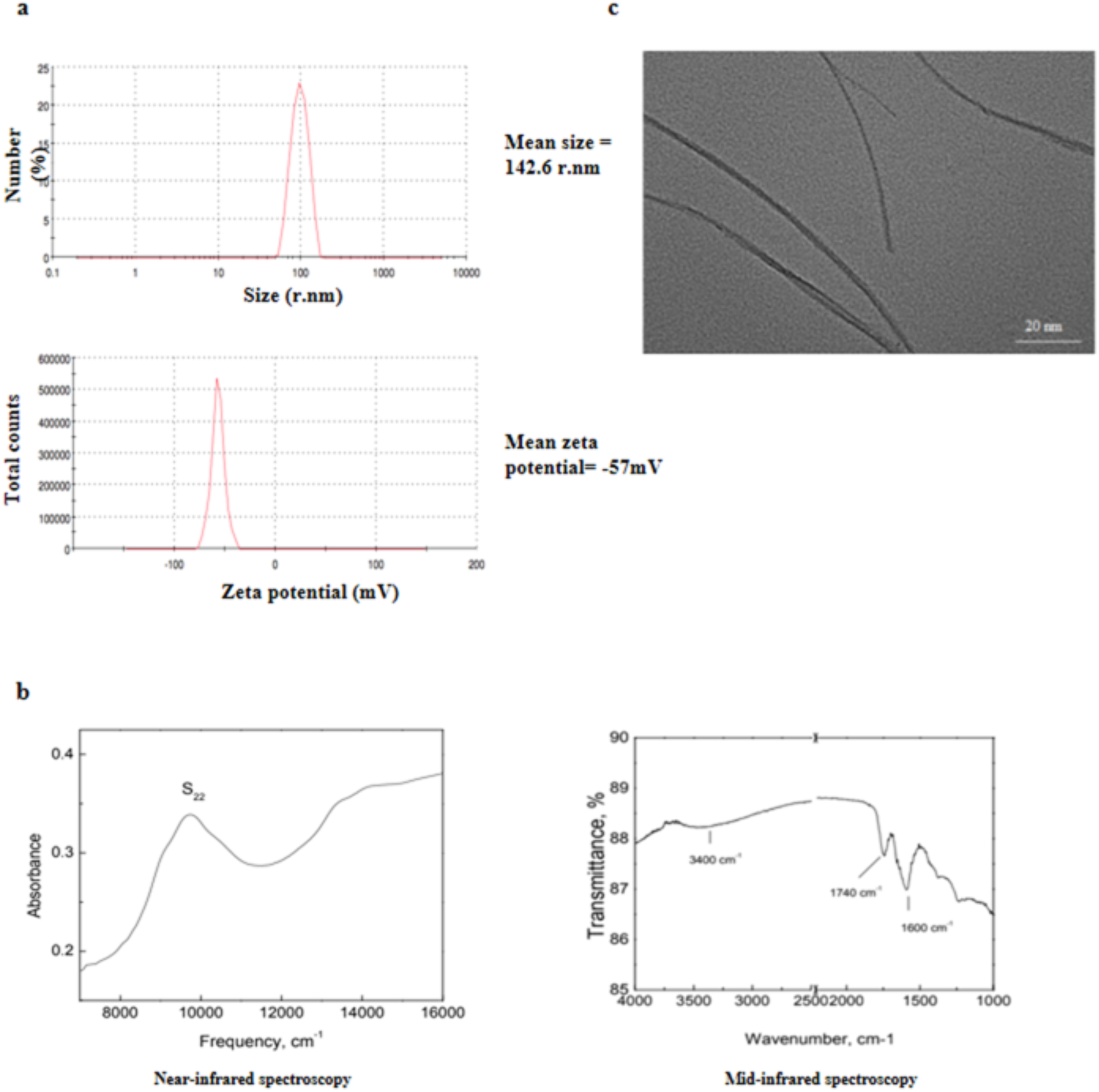
Characterization of AF-SWCNTs. Panel a shows the Zeta potential and Zeta size of AF-SWCNTs measured using Malvern Zetasizer. Panel b shows the mid-infrared and near-infrared spectroscopic results for AF-SWCNTs (Data from Sigma-Aldrich). Panel c shows the TEM image of strands of AF-SWCNTs.

**Supplementary Fig. 2.**
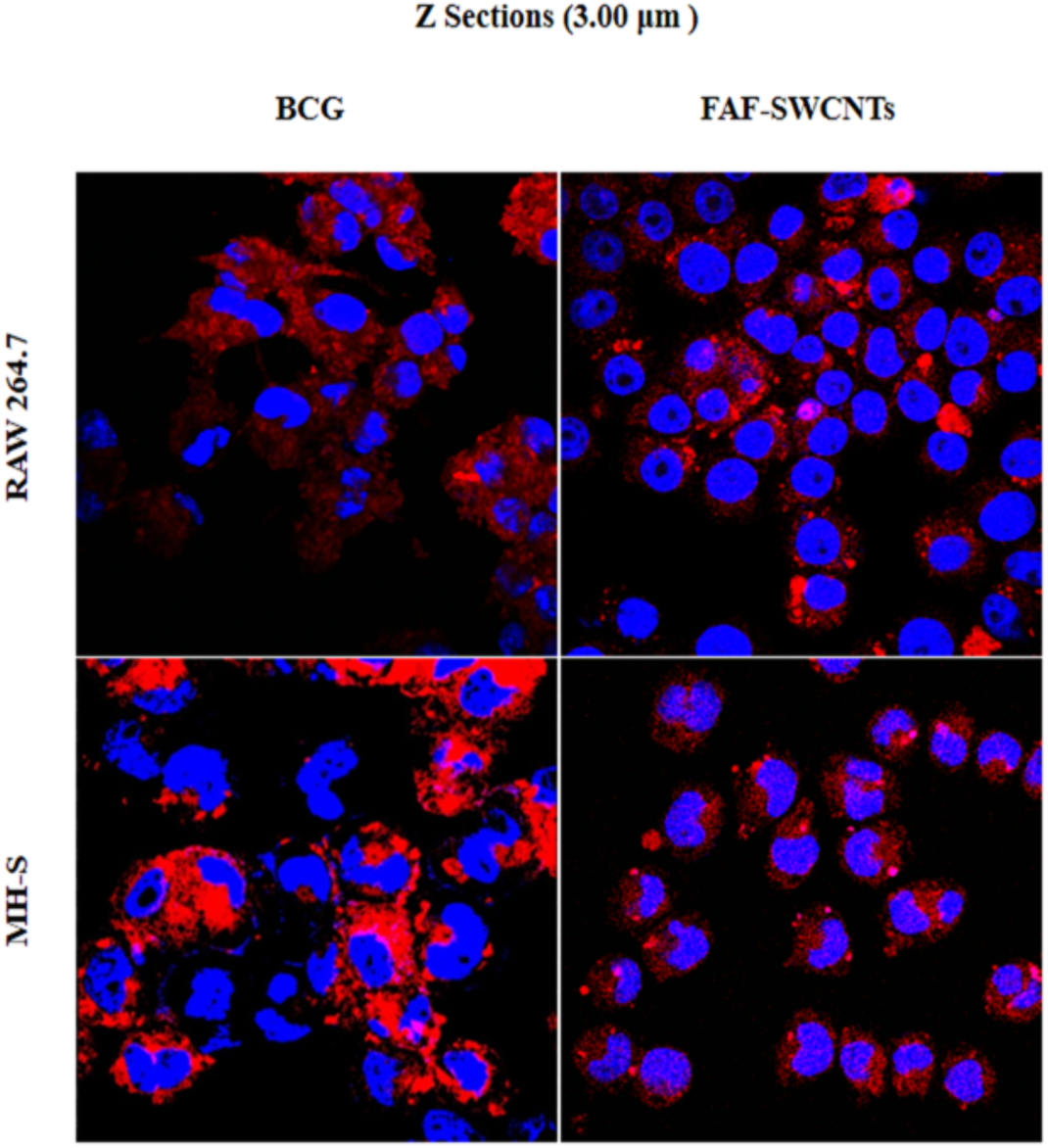
Z sectioning images showing uptake of DiI C-18 labelled BCG and FAF-SWCNTs by macrophages using confocal microscopy. RAW 264.7 and MH-S cells [0.1×10^6^/ml/well] were cultured on a coverslip with DiI C-18 labelled BCG (MOI 20:1) and FAF-SWCNTs (10 µg/ml) for 12 hours. After incubation, cells were washed with PBS, fixed using paraformaldehyde, stained nuclei using Hoechst 33342 dye and mounted on a glass slide to visualize the uptake of DiI C-18 labelled BCG and FAF-SWCNTs and performed Z sectioning using Confocal microscope. Confocal microscopy images showing internalization of labelled BCG by RAW 264.7 and MH-S are shown. In each image, micrographs represent Z sectioning images at 3 µm showing DiI C-18 labelled BCG uptake/FAF-SWCNTs uptake (red fluorescence) and nuclei are defined by blue stain of Hoechst 33342 dye. Magnification 100X in all cases.

